# An averaging strategy to reduce variability in target-decoy estimates of false discovery rate

**DOI:** 10.1101/440594

**Authors:** Uri Keich, Kaipo Tamura, William Stafford Noble

## Abstract

Decoy database search with target-decoy competition (TDC) provides an intuitive, easy-to-implement method for estimating the false discovery rate (FDR) associated with spectrum identifications from shotgun proteomics data. However, the procedure can yield different results for a fixed dataset analyzed with different decoy databases, and this decoy-induced variability is particularly problematic for smaller FDR thresholds, datasets or databases. In such cases, the nominal FDR might be 1% but the true proportion of false discoveries might be 10%. The averaged TDC protocol combats this problem by exploiting multiple independently shuffled decoy databases to provide an FDR estimate with reduced variability. We provide a tutorial introduction to aTDC, describe an improved variant of the protocol that offers increased statistical power, and discuss how to deploy aTDC in practice using the Crux software toolkit.

## Introduction

Tandem mass spectrometry (MS/MS) can be thought of as a high-throughput technique for generating hypotheses, wherein complex biological samples are analyzed to yield potential insights into their protein contents. Accordingly, taking action on the basis of MS/MS results requires a method for assigning statistical confidence to these hypotheses. Our willingness to perform laborious confirmatory experiments on a set of detected peptides, for example, will be higher if we believe that the proportion of false discoveries in the peptide list is at most 1%.

In the scenario we focus on here—bottom-up data-dependent acquisition tandem mass spectrometry—the most basic type of hypothesis concerns the generation of an observed fragmentation spectrum by a particular charged peptide species. In principle, the corresponding peptide-spectrum match (PSM) can either be correct or incorrect, depending upon whether the specified peptide is responsible for generating the observed spectrum.

Accordingly, an extensive literature focuses on the problem of assigning statistical confidence estimates to individual PSMs or to collections of PSMs. This literature is quite complex, involving diverse methods such as expectation-maximization, ^1^ parametric fitting of PSM scores to Poisson, ^2^ exponential,^3;4^ or Gumbel distributions,^5;6^ nonparametric logistic regression, ^7^ various types of dynamic programming procedures, ^8–10^ as well as machine learning post-processors, such as linear discriminant analysis ^1^ or support vector machines. ^11–13^

Despite this dizzying array of techniques, by far the most widely used method for assigning statistical confidence estimates to PSMs is quite straightforward and easy to understand. The method, called “target-decoy competition” (TDC), involves assigning peptides to spectra by searching the spectra against a database that contains a combination of real (“target”) peptide sequences and reversed or shuffled (“decoy”) peptides. ^14^ The decoys, which by definition represent incorrect hypotheses, provide a simple null model. Accordingly, for every decoy PSM that we observe, we estimate that one of the target PSMs is also incorrect. Thus, the TDC protocol simply involves ranking PSMs by their scores and then counting the number of targets and decoys observed at a specified score threshold.

Although TDC is employed routinely in many shotgun proteomics studies, the fact that the FDR estimates that TDC produces can exhibit high variability is not widely appreciated. In practical terms, this means that a set of PSMs with a nominal FDR of 1% might actually contain, say, 5% incorrect PSMs. We begin by illustrating this phenomenon, demonstrating that the level of variability increases as the size of the dataset or the size of the peptide database decreases. We then explain how to apply our previously described “average TDC” (aTDC) procedure to reduce the variability in the estimated FDR by searching multiple, independently shuffled decoy databases. ^15^ Furthermore, we introduce an improved version of the aTDC protocol that yields better statistical power (i.e., more accepted PSMs at a fixed FDR threshold), especially for low FDR thresholds. Finally, we provide a detailed protocol for applying this improved aTDC procedure using the Crux mass spectrometry toolkit. ^16^ Note that, in keeping with the tutorial nature of this paper, we have opted to move the Methods section, which includes technical details, to the end and to begin the Results section with intuitive descriptions of TDC and aTDC.

## Results

### Target-decoy competition yields confidence estimates but sacrifices some identifications in the process

Explaining how the aTDC protocol works requires first carefully explaining the TDC protocol. Furthermore, to make the aTDC explanation easier later, we divide our description of the TDC protocol into three steps, even though in practice the first two steps are typically carried out jointly by searching a concatenated database of targets and decoys.

In the first step, a given collection of spectra (a dataset) is searched against two peptide databases: the target database of interest and a corresponding decoy database. We assume that each target peptide has a corresponding decoy peptide, where the decoy is a shuffled or reversed version of the target. The database search procedure can be carried out by any search engine (reviewed in Verheggen *et al*. ^17^). For each spectrum, this search identifies a best-scoring target peptide and a best-scoring decoy peptide (Figure 1).

**Figure 1:**
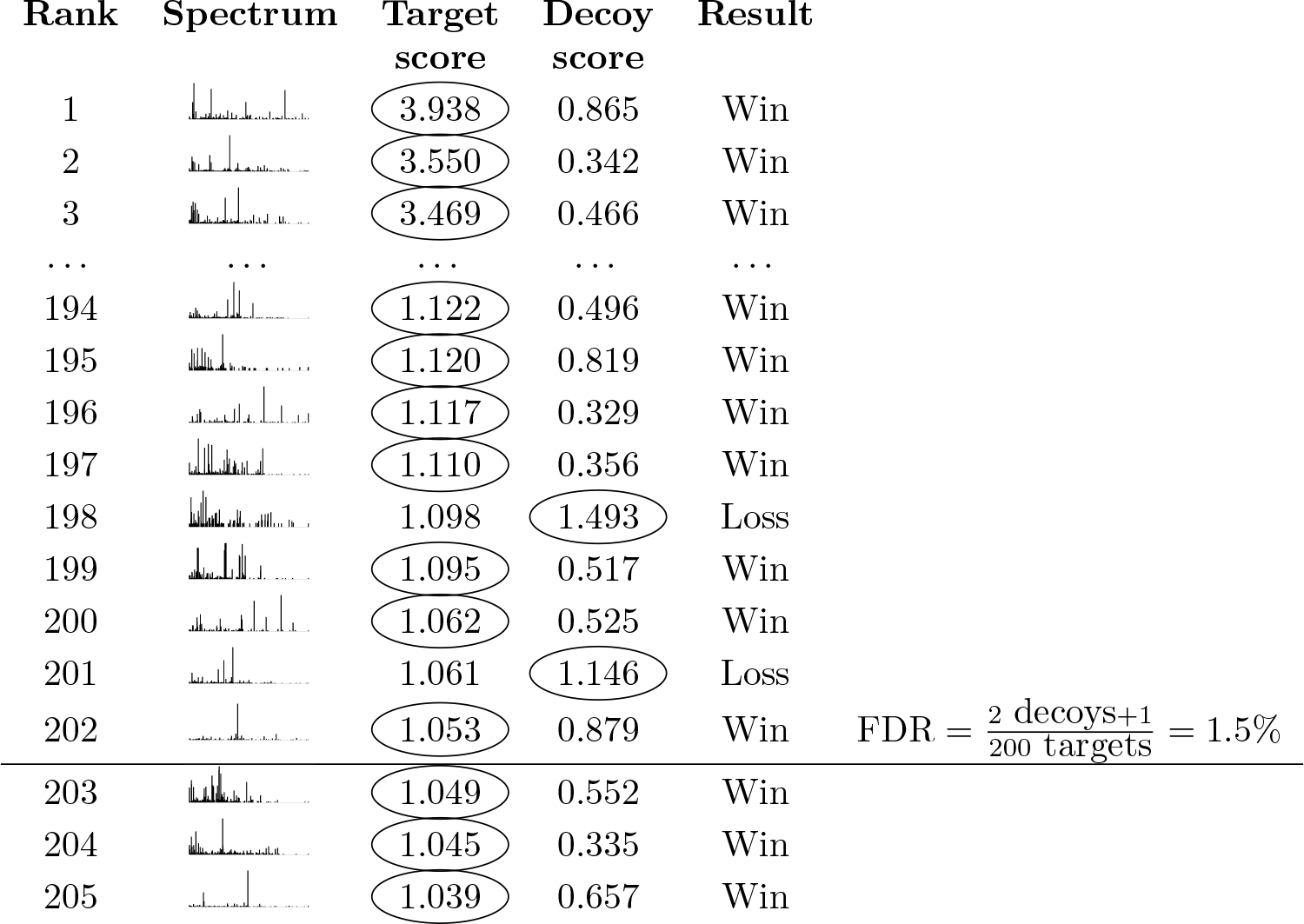
Target-decoy competition. For each spectrum, a database search procedure identifies the top-scoring target peptide and the top-scoring decoy peptide. The target and decoy then compete such that the better of the two matches is assigned to the spectrum. The FDR among target PSMs at a given score threshold *ρ* is then estimated via Equation 1. Including “+1” in the numerator makes this the TDC^+^ protocol.

In the second step, for each spectrum we compare the scores assigned to the target and the decoy, and we designate the PSM with the higher score as the winner, assuming that higher scores indicate better matches. This is the “competition” step of TDC.

Finally, in step three, we sort the winning PSMs by score and set a score threshold *ρ*. We will refer to PSMs with scores that exceed our specified threshold as “accepted” PSMs. The key idea of TDC is that, for each decoy PSMs we accept, we estimate that we have also accepted an incorrect target PSM. Accordingly, TDC estimates the FDR associated with the set of accepted target PSMs as

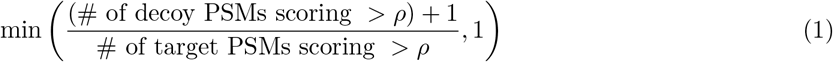

Two features of Equation 1 require further explanation. First, the “min” operation in Equation 1 is only included to account for the (hopefully rare) case where the number of decoys is at least as large as the number of targets at a specified threshold. Without the min operation, the estimated FDR could exceed 100%. Second, the numerator (the number of decoys scoring > *ρ*) has a “+ 1” added to it. This +1 correction was not included in the original description of TDC. ^14^ Accordingly, we will refer to the TDC procedure that incorporates this +1 correction as “TDC^+^,” in order to distinguish it from the original TDC protocol. The need for this type of correction was proved by Barber and Candés in the context of linear regression^18^ (see their “knockoff^+^” procedure) and by He *et al.* in the context of mass spectrometry (see their Equation 25). ^19^ Subsequently, Levitsky *et al.* provided a more intuitive explanation of the need for the +1 correction in shotgun proteomics. ^20^

A key feature, and unavoidable drawback, of the TDC (or TDC^+^) protocol is that the competition has the undesirable side effect of randomly discarding some high-scoring target PSMs. For example, in Figure 1, spectra 198 and 201 matched a target peptide with scores greater than the specified threshold. Unfortunately, however, these high-scoring target PSMs got unlucky: they happened to be outscored by a random decoy peptide. In practice, the rate at which this random loss of high-scoring PSMs occurs is typically low, but the magnitude of the problem increases as the size of the peptide database grows, as well with a more permissive FDR threshold.

### Using shuffled decoys leads to variation in the list of identified spectra

It should be clear, given the above description, that the TDC procedures only provide an estimate, not an exact count, of the number of incorrect accepted target PSMs. This is to be expected but leaves open the question of how variable the estimate is.

One simple way to address this question is to re-shuffle the decoy database and then repeat the TDC procedure. We carried out such an experiment, using a single mass spectrometry run selected from the Kim *et al.* “draft human proteome” dataset^21^ and searched against the human proteome using the widely used XCorr score function^22^ followed by TDC^+^. Ten searches against different decoy databases are included. The results (Figure 2A) show a surprisingly high degree of variability. The first time we ran the search we accepted 4987 target PSMs at a 1% FDR threshold, but after nine additional searches the count of accepted PSMs ranged from a minimum of 4757 to a maximum of 4987 (by chance, our first random shuffle provided the greatest number of discoveries). We quantify the observed variability by computing the percentage difference between the maximum and minimum, relative to the mean of the maximum and minimum. In this case, this variability is 4.7%. The percent variability is lower at less strict FDR thresholds (1.6% at a 5% FDR threshold, and 3.2% at 10% FDR).

**Figure 2:**
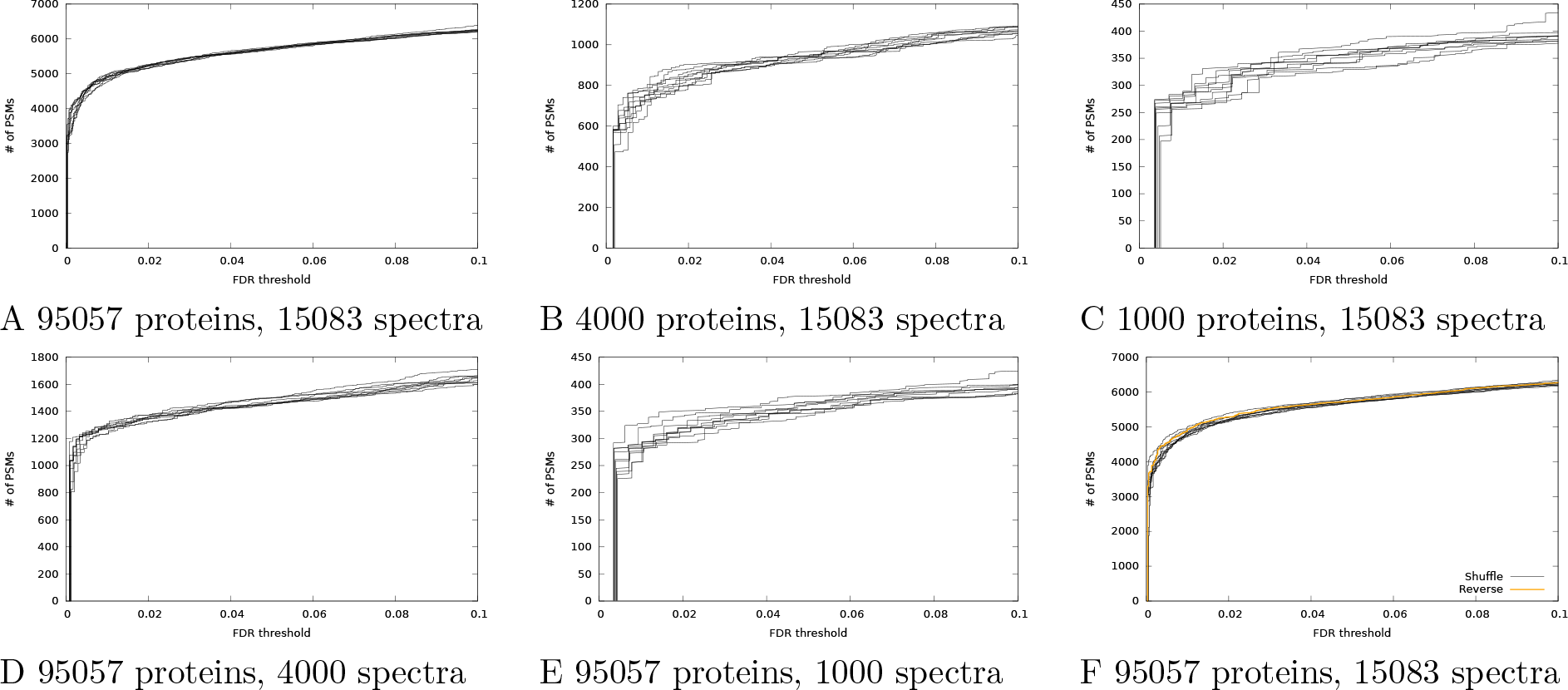
Variation in discoveries from TDC^+^. (A) The figure plots the number of accepted PSMs as a function of FDR threshold for the Kim dataset, searched using Tide with the XCorr score function. Results from ten searches against different decoy databases are shown. (B-C) Same as panel A, but after randomly downsampling the database to contain fewer proteins. (D-E) Same as panel A, but after randomly down-sampling the dataset to contain fewer spectra. (F) Same as panel A, but also including a line corresponding to TDC^+^ with reversed peptide decoys.

A similar trend can be observed when we vary either the number of spectra included in our search (Figure 2B-C) or the number of proteins in the sequence database (Figure 2D-E). For example, when we decrease the size of the database from 95,057 to 4000 proteins, the percentage difference between the minimum and maximum numbers of accepted target PSMs increases from 1.6% to 5.6% at a fixed FDR threshold of 5%. A similar trend at a 5% FDR threshold is seen when we decrease the number of spectra from 15,083 (1.6% variability) to 4000 (3.7% variability) or 1000 spectra (10.0%).

### Reversing the decoys does not eliminate the variability

At this point, many readers may be thinking to themselves, “Luckily, I am immune to this variability problem because I use reversed peptide decoys rather than shuffled peptide decoys.” Unfortunately, reversal does not solve this problem; it merely makes the problem harder to see.

To understand why this is so, consider a “fixed permutation” strategy for generating decoys. One could imagine generating decoy peptides by taking each target peptide and permuting it according to some fixed rule; e.g., swap the amino acids in positions 1 and 3, then positions 2 and 4, etc. Using such an approach, and armed with ten different sets of permutation rules, one could generate results similar to those in Figure 2. In this setting, the reversal scheme would represent just one, arbitrary choice of fixed permutation.

We repeated the search of the Kim dataset but using a set of reversed peptide decoys. As expected, the resulting curve (Figure 2F) falls into the middle of the curves generated by shuffled decoys.

### Average target-decoy competition leads to reduced variability

The aTDC protocol was designed to reduce variability in decoy-based FDR estimates without sacrificing statistical power. The method works by creating multiple decoy databases, each one equal in size to the target database. Consequently, if we run aTDC with, say, three decoy databases, then for each observed spectrum we obtain one target score and three decoy scores (Figure 3). The key idea behind aTDC is to estimate that each decoy PSM that wins the target-decoy competition corresponds to 1/3 of an incorrect target PSM. Consequently, we only lose one target for every three decoys that win the competition. The final FDR estimation is still performed using Equation 1, where the “+1” in the denominator can be included (for aTDC^+^) or not (for aTDC). All experiments reported here use aTDC^+^ or a new variant thereof, introduced below.

**Figure 3:**
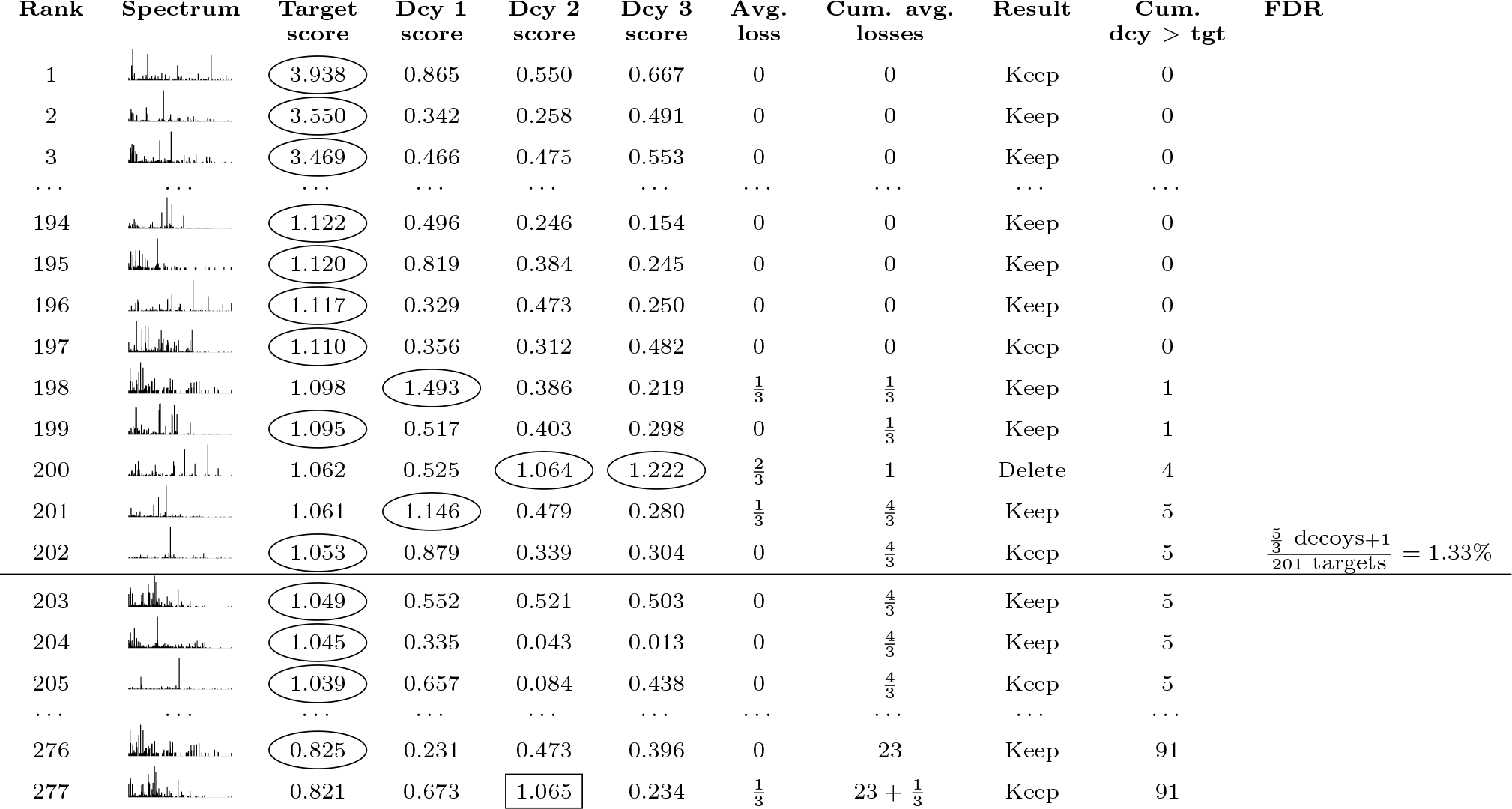
Average target-decoy competition. The procedure is similar to TDC (Figure 1), except that each target competes against multiple decoys and losses are counted fractionally. A target is eliminated only when the rounded fractional count of losses reaches the next integer value. As in TDC^+^, FDR for aTDC^+^ is estimated via Equation 1. Note that decoys for lower-ranking targets that receive high scores (e.g., decoy 2 in line 277) can contribute to the cumulative decoy count for lines above them in the list.

The aTDC^+^ protocol yields FDR estimates that are consistent with TDC^+^ but that exhibit markedly less variability. To illustrate the effect, we re-ran the search of the Kim dataset 10 times with aTDC^+^, each time using 10 decoy databases. The resulting curves overlay the corresponding curves from TDC^+^ but exhibit much lower variability (Figure 4). At 5% FDR, the percentage change between the minimum and maximum number of accepted target PSMs reduces from 1.6% for TDC^+^ to 0.52% for aTDC^+^. We also note a slight increase in power of aTDC^+^ relative to TDC^+^ in Figure 4. This effect arises because aTDC selects which target PSMs to filter based on the number of times a given target won the target-decoy competition, and hence it is partially calibrating the score (see Keich and Noble ^15^ for details).

**Figure 4:**
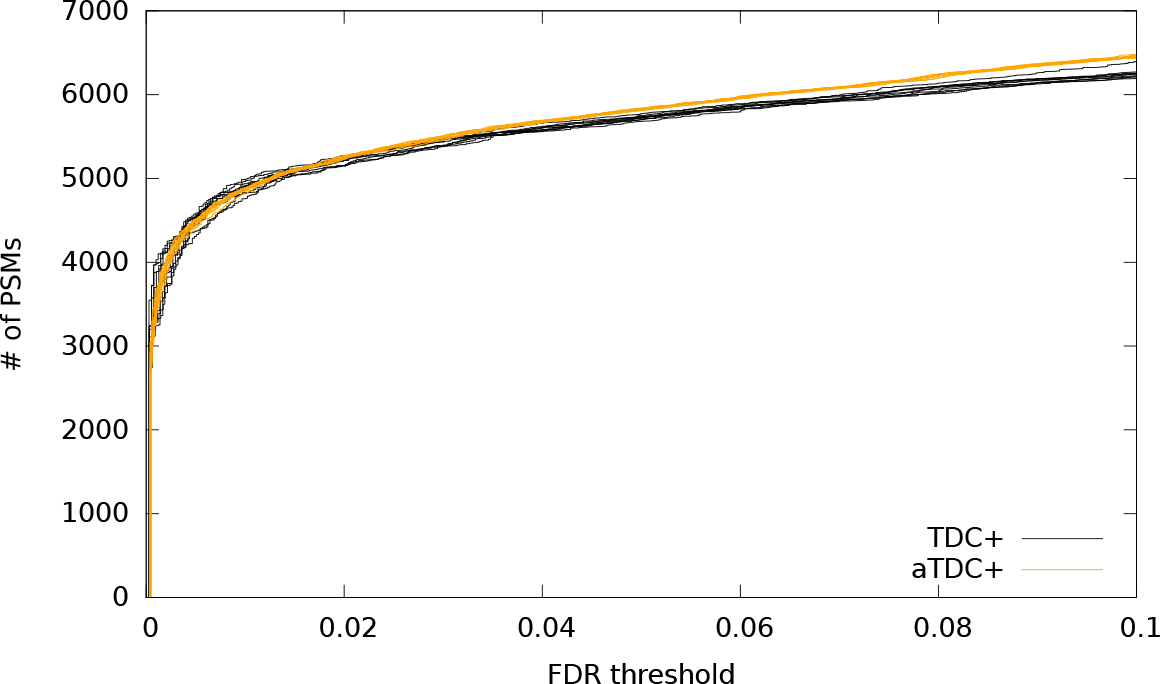
Average target-decoy competition leads to reduced variability. The figure plots, for the human dataset, the number of PSMs accepted as a function of FDR threshold for 10 runs of TDC^+^ and 10 runs of aTDC^+^. Each aTDC^+^ run is computed with respect to 10 decoy databases.

### The improved version of aTDC^+^ is less conservative than TDC^+^ for low FDR thresholds

One feature of the aTDC protocol, as originally described, ^15^ is that it avoids TDC’s small liberal bias (i.e., underestimation of FDR), which has been observed previously for low FDR thresholds.^23^ However, if we correct this bias by switching to TDC^+^ and aTDC^+^, then we observe that aTDC^+^ exhibits a pronounced conservative bias for the same low FDR thresholds. The effect can be seen, for example, in the shift of aTDC^+^ curves, relative to the TDC^+^ curves, at very low FDR thresholds (< 0.01) when we search the Kim dataset using a database of 1000 proteins (Figure 5A).

**Figure 5:**
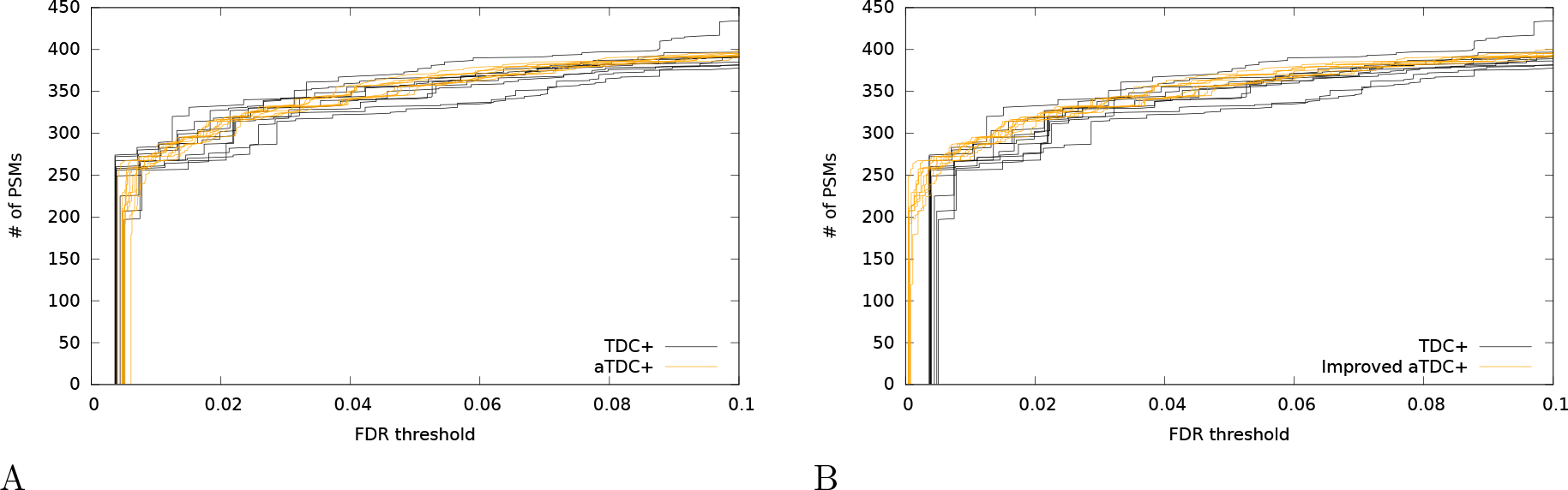
Improved version of average target-decoy competition reduces conservative bias for low FDR thresholds. (A) The figure plots, for the Kim dataset applied to a database of 1000 proteins, the number of PSMs accepted as a function of FDR threshold for 10 runs of TDC^+^ and 10 runs of aTDC^+^. (B) Same as panel A, but using the improved aTDC^+^ procedure (called 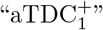 in the main text). In both panels, each run of aTDC^+^ or 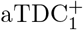 is computed with respect to 10 decoy databases.

To combat this problem, we have devised an improved version of aTDC^+^, denoted as 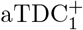, that provides better statistical power than TDC^+^ while still empirically controlling the FDR. As noted above, the only difference between aTDC^+^ and 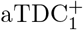 lies in the correction factor that we add to the number of decoy discoveries at the *i*th PSM in the ranked list (i.e., the *i*th row in Figure 3). In aTDC^+^, we use +1, whereas in 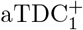 we essentially use the average number of decoy scores that fall between the target scores with ranks *i* and *i* − 1.

To motivate this change we go back to the explanation in Levitsky *et al.* ^20^ of why we need to add the +1 term in (1) in the first place. As shown in Theorem 1 of He *et al.*,^19^ the FDR estimate that does not involve the +1 term is in fact a *conservative* estimate of the FDR for each fixed *t*. The intuitive reason it is still liberally biased in the context of TDC is that we select our cutoff score as the smallest score threshold for which the estimated FDR is below the desired level. This implies that we conveniently select our threshold so that the next largest score corresponds to a decoy discovery which we have just excluded. This exclusion creates a slight liberal bias which the +1 term is correcting: essentially we are allowing for the subsequent decoy discovery.

In the context of aTDC we have multiple decoys, and we average their numbers of discoveries. Therefore, rather than blindly adding +1 to the number of decoy discoveries at the *i*th PSM in our observed ranked list, we postulate that we do not need to add more than the average number of decoys we observed with scores between the scores associated with the *i*th and *i* + 1th PSM. Note that this is equivalent to the difference between the *i*th and *i* − 1th entries of the “Cum dcy > tg” column in Figure 3. Intuitively, this number is an estimated bound on the expected number of decoy discoveries we exclude when setting the score threshold as we do.

Repeating the analysis of the Kim dataset with a database of 1000 proteins, we observe that the average number of PSMs at 1% FDR improves from 271 for aTDC^+^ to 284 for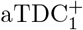. Supplementary Note 1 provides extensive evidence that the resulting FDR estimates are unbiased.

To facilitate adoption of 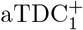 by the research community, we have implemented the protocol within the Crux mass spectrometry toolkit (http://crux.ms).^16^ The procedure consists of creating a peptide index containing multiple decoy databases, searching with the index using the Tide search engine, and then post-processing the resulting target and decoy PSMs using the assign-confidence command in Crux. A detailed protocol, with supplementary input and output files provided, is provided in Figure 6.

**Figure 6:**
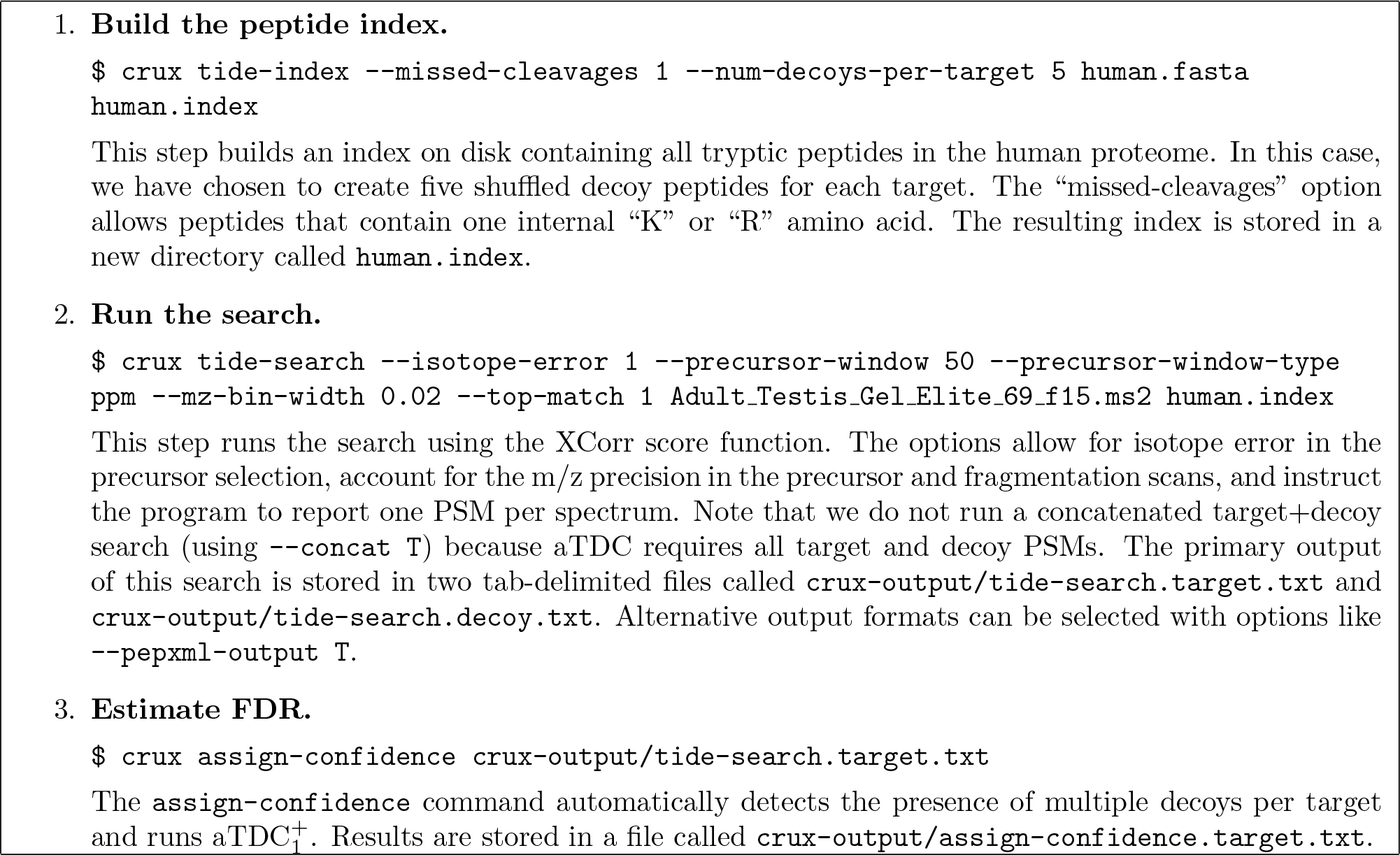
Performing average target-decoy competition using Crux. The procedure assumes that the Crux software is already installed on your computer and that files containing the spectra and the human proteome database reside in the current directory. These files are available in Supplementary File S2. Further documentation about the Crux commands used above is available at http://crux.ms.

### Selecting the number of decoy databases

In practice, carrying out the protocol in Figure 6 requires that the user decide how many decoy databases to create in the very first step. Unfortunately, it is not always obvious how to select this number. Here, the trade-off is between reducing the variability of the FDR estimate versus the computational cost associated with searching multiple decoy databases. Clearly, this trade-off is something that cannot be decided on the basis of statistical theory, because it depends upon the time and resources available to the analyst.

To assist users in assessing this time-vs-variability trade-off, we have performed a systematic study that estimates how 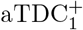 reduces variability as a function of the number of decoy databases, the number of spectra, the size of the peptide database, and the FDR threshold. For this study, we use ten randomly selected runs from the Kim dataset. Note that throughout the preceding exposition we have used as our measure of variability the percentage difference between the minimum and maximum number of accepted target PSMs. We selected this measure because it is intuitive and easy to understand. However, for our empirical study, we have instead opted to use the empirical standard deviation of the estimated FDR, since standard deviation is a more robust statistical measure.

The results of this empirical survey (Figure 7) show consistent trends across different FDR thresholds and varying sizes of databases and datasets. In particular, we observe a rapid decrease in decoy-induced variability when using even just a single additional decoy database. In nearly every case, the standard deviation continues to decrease as the number of decoys increases to 3, 5, 10 and 25. Not surprisingly, we also observe a diminishing returns property, such that each additional decoy database yields a smaller reduction in standard deviation as the total number of decoys increases. Given the trends in Figure 7, and taking into account the logarithmic y-axis, we suggest that a reasonable cost-benefit trade-off might be to employ five decoy databases. This represents a three-fold increase in computational cost (searching one target plus five decoy databases compared to searching one target plus one decoy) while eliminating a large proportion of the decoy-induced variability.

**Figure 7:**
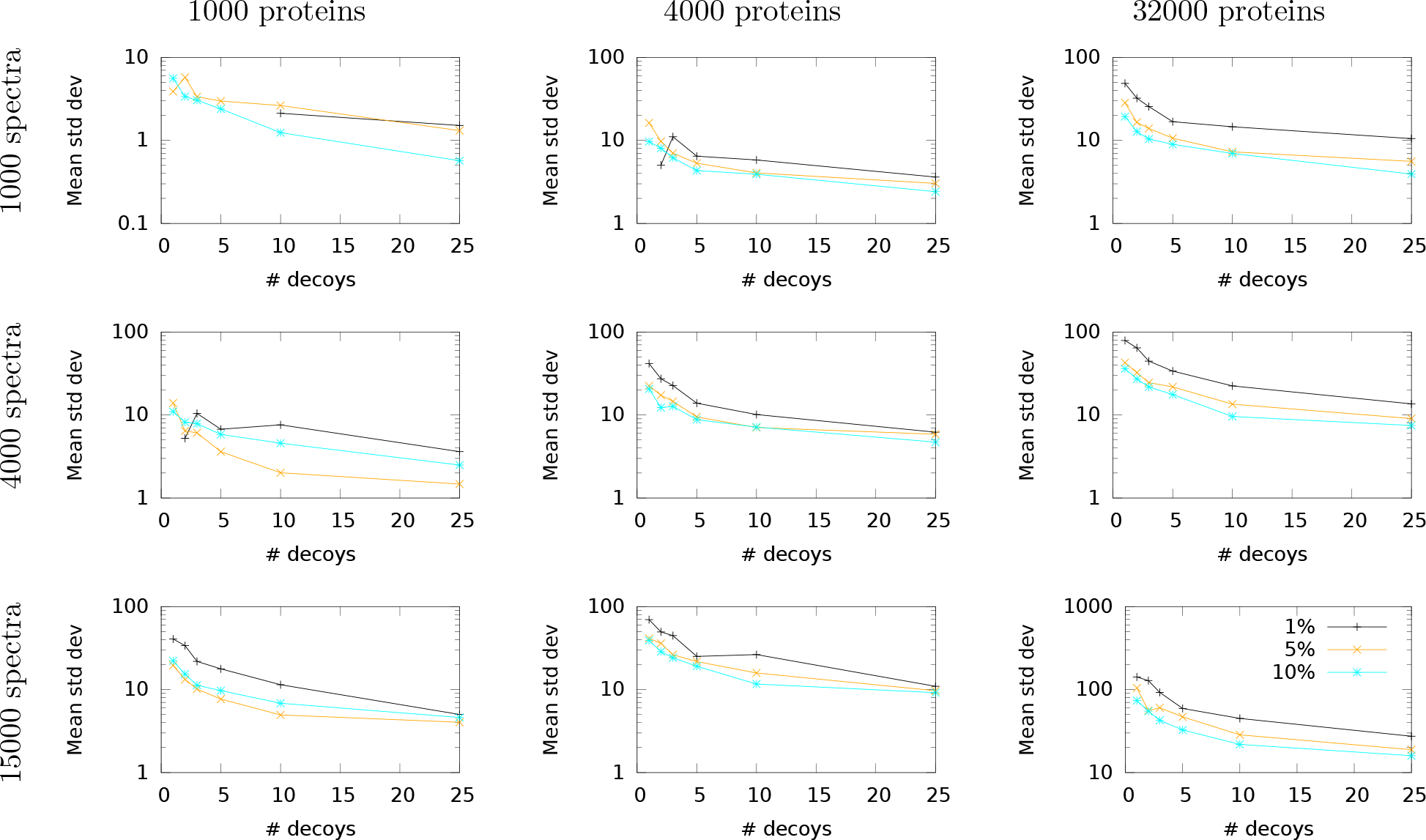
Empirical study of variability of FDR estimates. Each panel plots the standard deviation in the number of accepted target PSMs, averaged over ten mass spectrometry runs from the Kim dataset, as a function of the number of decoys used for 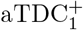. Each line corresponds to a different FDR threshold. All standard deviations are calculated over ten repetitions of either TDC^+^ (for 1 decoy) or 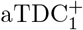 (for >1 decoy). Missing points correspond to cases that yielded a standard deviation of zero across all ten runs.

## Discussion

The average target-decoy competition protocol provides a straightforward and unbiased method for decoy-based estimation of the FDR among a given set of peptide-spectrum matches, while providing the user a means of reducing decoy-induced variability in the resulting estimate. In addition to explaining how the method works, we have provided an open source software implementation and an improved algorithm 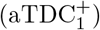 that yields better statistical power at low FDR thresholds.

Some readers may wonder whether aTDC is more complex than it needs to be. In particular, perhaps the most natural approach to reducing decoy-induced variability is simply to increase the size of the decoy database: rather than shuffling each target peptide once, we could shuffle each peptide, say, 10 times, yielding a decoy database that is 10 times larger than the target database. This change requires a simple adjustment to the TDC protocol, such that each time we see an accepted decoy PSM, we estimate that it corresponds to 1/10th of an incorrect target PSM. This simple approach does indeed reduce the empirical variability in the estimated FDRs. However, particularly for a well-calibrated score function, expanding the decoy database in this way also leads to a decrease in statistical power; i.e., for a fixed FDR threshold, using a larger decoy database yields, on average, fewer accepted target PSMs than using a smaller decoy database. The source of this loss in statistical power is the “competition” component of TDC. As the size of the decoy database increases, each decoy has more chances to randomly achieve a high score. Hence, the rate at which high-scoring targets are eliminated due to competition with targets increases. The 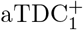 protocol avoids this trade-off, delivering reduced variability along with statistical power that is equal to or better than TDC^+^.

So far we have discussed the variability induced by the selection of decoy peptides, but this is only part of the story. Additional variability is produced by the “draw” of the spectrum set. Specifically, that set is a combination of native spectra generated by peptides in the database and foreign spectra generated by molecules not in the database. The relative proportions of each of these two components as well as the noise in the generation of each native spectrum inject random effects into the experiment. This randomness is independent of the decoy database but could still have a significant impact on FDR estimation and is not something we can eliminate.

The 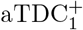 protocol uses multiple decoy databases to reduce decoy-induced variability, but we have also previously described an alternative use of multiple decoy databases to instead improve statistical power. The “progressive calibration” procedure provides a method for increasing the number of PSMs accepted at a specified FDR threshold by calibrating PSM scores using a collection of matched decoy scores. ^15^ Like 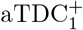, progressive calibration’s utility increases as the number of decoys increases, with diminishing returns as each successive set of decoys is added. Thus, a user with sufficient resources to generate a fixed number of decoy databases can decide how to apportion those databases to achieve both reduced variability and improved power. In future, we plan to provide an implementation of progressive calibration in Crux, to complement the 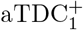 implementation.

We have also identified several other avenues for future work. For example, it is not yet clear how to combine 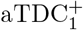 with a post-processor like Percolator. ^13^ Percolator employs decoys for two purposes: first, to train a classifier to discriminate between targets and decoys, and second, to estimate FDR. A cross-validation scheme is necessary in order to ensure that the same PSMs are not used in training the classifier and in FDR estimation. ^24^ Fitting a collection of decoy databases into this cross-validation scheme while ensuring valid FDR estimation is non-trivial. Another direction for future work is to extend the averaging idea to FDR estimation at the peptide and protein levels.

**Table 1.** Variables and their definitions.

| Variable | Definition |
| --- | --- |
| $\Sigma$ | set of all spectra (observed) |
| $n$ | number of spectra (observed) |
| $\mathcal{D}^t$ | target database of peptides (user determined) |
| $\mathcal{D}_i^d$ | decoy database of peptides (user determined) |
| $n_d$ | number of competing decoy database (user determined) |
| $\rho$ | score threshold (user determined) |
| $D(\rho)$ | number of decoy discoveries (at level $\rho$) in a search of the concatenated database $\mathcal{D}^t \cup \mathcal{D}^d$ (observed) |
| $T(\rho)$ | number of target discoveries (at level $\rho$) in a search of the concatenated database $\mathcal{D}^t \cup \mathcal{D}^d$ (observed) |
| $\sigma$ | single spectrum (observed) |
| $w(\sigma)$ | score of the best match of $\sigma$ in the target database (observed) |

## Methods

### Target-decoy competition

In our setting, a decoy-based FDR controlling procedure takes as input a list of target PSMs produced by searching a set ∑ of spectra against a database 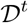 of real (“target”) peptides and a corresponding list of decoy PSMs produced by searching the same spectra against a database 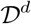 of decoy peptides. The decoy peptides are created by the user and can be either shuffled or reversed versions of the targets. For any score threshold *ρ*, the TDC procedure defines its list *T* (*ρ*) of discoveries as all target PSMs with score ≥*ρ* that outscore their corresponding decoy competition. Note that this definition is equivalent to saying that these PSMs remain discoveries in a search of a concatenated database 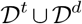. Note also that we assume the score is defined such that larger values correspond to better matches. Denoting by *D* (*ρ*) the number of decoy discoveries at score level *ρ* in the concatenated search, TDC estimates the FDR in its target discovery list, at level *ρ*, as

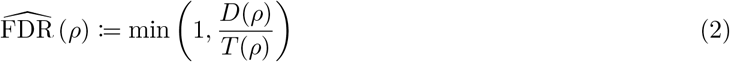

where the min() operation is included to handle the rare case where the number of decoy discoveries exceeds the number of target discoveries at the specified threshold. Note that Equation 2 is equivalent to Equation 1.

We also define a variant of TDC, called “TDC^+^,” that incorporates a small theory-mandated correction that is particularly important for small sample sizes (more on that in the Results section). TDC^+^ is identical to TDC except that the FDR is estimated as

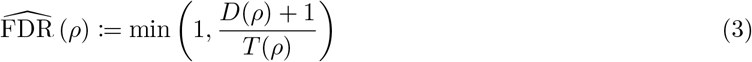

Both TDC and TDC^+^ set the discovery score cutoff to be the smallest observed score *ρ* for which the corresponding estimated FDR is still below the desired FDR level.

### Average target-decoy competition

The average TDC procedure is similar to TDC except that the spectra ∑ are searched against the target database 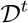 and *n* independently generated (typically, shuffled) decoy databases 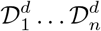. The procedure then consists of the following steps:

- Sort the set of optimal target PSM scores, {*w* (σ) : σ ∈ ∑} in decreasing order and denote them by *ρi*.
- For every decoy database 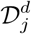, apply TDC to the target database 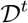 and 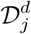 and note the corresponding number of target, *T_j_* (*ρi*), and decoy, *D_j_* (*ρi*), discoveries at level *ρi*.
- Use the above TDC data to compute the average number of target and decoy discoveries at score level 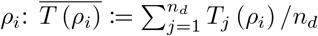 and 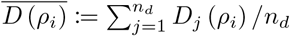.
- Initiate *T_c_*, the cumulative number of target discoveries, to 0 and set 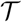, the target discovery indicator, to a logical vector of size *n* (initiated to all TRUE)
- From the largest (best) target score to the smallest (*i* = 1 : *n*) do:

— if 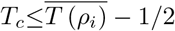 set *T_c_* = *T_c_* + 1 and skip to next *i*
— else

* Consider all current target discoveries with score ≥ *ρi* (i.e., all *j* ≤ *i* for which 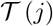 is currently TRUE).
* Restricting attention to those discoveries in the above list that lost the most decoy competitions, choose the one with the smallest score.
* Set the 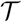 entry for that chosen target PSM to false (i.e., remove it from the current list of target discoveries) and continue to next *i*. Please refer to Keich and Noble ^15^ to see how the last step can be efficiently implemented.
- Set the vector *T* that yields the number of discoveries at level *ρi* to the cumulative sum of the vector 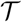
- Set 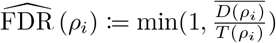 for all *i*
- Set the cutoff score to be the smallest *ρi* for which 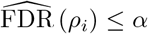, where *α* is the selected FDR threshold

Analogous to TDC and TDC^+^, we also define a variant of aTDC (aTDC^+^) that replaces the FDR estimation in the final step with 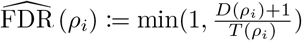

### Improved variant of average target-decoy competition

We introduce 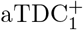, a more powerful version of aTDC^+^, where we estimate the FDR in the last step as,

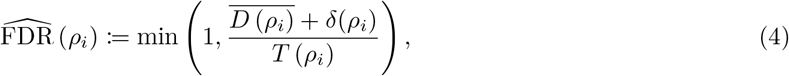

where 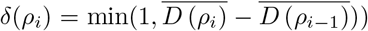. The motivation for this modification is given in Results; here we only note that because *δ*(*ρi*) ≤ 1, the FDR estimated in 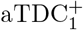 is generally no larger than the one estimated in aTDC^+^, and hence 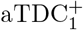 will report at least as many discoveries as aTDC^+^.

### Datasets and analysis

All of the primary analyses are performed using a single run selected from the “draft human proteome” of Kim *et al.* ^21^ This file (Adult_Testis_Gel_Elite_69_f15) contains 15083 spectra.

All searches are carried out with respect to the Uniprot human reference proteome, downloaded on 12 July 2018 and including multiple isoforms per protein. This database contains 95,057 protein sequences.

Peptide indices are created using tide-index, allowing for clipping of N-terminal methionines (clip-nterm-methionine=T), at most one missed cleavage (missed-cleavages=1), a variable N-term peptide modification of 42.0367 Da (nterm-peptide-mods-spec=1X+42.0367), and allowing duplicates in the decoy database (allow-dups=T). This yields a peptide index containing 3,697,160 distinct target peptides.

All searches are performed using Tide^25^ with the default XCorr score function. Additional search parameters include a precursor window of 50 ppm (precursor-window-type=ppm precursor-window=50), an m/z bin width of 0.02 Da (mz-bin-width=0.02), one isotope error (isotope-error=1), and reporting a single match per spectrum (top-match=1).

## Supplemental files

- **Supplemental File S1: Empirical analysis of the power and FDR control of** 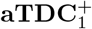
- **Supplemental File S2: Protocol for using** 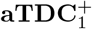 **in Crux**. The file “protocol.zip” contains all input files necessary to carry out the protocol outlined in Figure 6. Also included are the log files produced by all three Crux commands, as well as the output files produced by `tide-index` and `assign-confidence`. This file is too large for upload so can be found at https://noble.gs.washington.edu/proj/atdc/protocol.zip.

